# PhytoScan3D: an open-source Python pipeline for batch extraction of phenotypic traits from 3D point cloud files generated by multispectral plant phenotyping sensors

**DOI:** 10.64898/2026.06.01.729298

**Authors:** Mallikarjuna Rao Kovi, Antonio Candea Leite, Morten lillemo

## Abstract

High-throughput 3D multispectral plant phenotyping platforms generate large volumes of point cloud files, but trait extraction is typically performed by sensor-bundled software whose internal algorithms are not publicly documented, which limits reproducibility and integration into custom research pipelines. Here we present PhytoScan3D, an open-source Python pipeline that extracts morphological and spectral phenotypic traits, spanning plant height, 3D leaf area, digital biomass, convex hull volume, leaf inclination, canopy geometry, NDVI, hue, and vegetation indices, from both PLY and PCD point cloud files generated by Phenospex PlantEye F500 and F600 sensors, and is portable to point clouds from any acquisition platform. PhytoScan3D was validated against HortControl (PhenoSpex) ground-truth measurements on 936 barley (*Hordeum vulgare*) pot-date observations from the growth chamber trial (20 Norwegian cultivars, 12 scan dates, Septemenr 2025 to January 2026), achieving Pearson r = 0.913 to 0.999 and ratio approximately 1.000 for Plant Height Max, 3D Leaf Area, and NDVI Average. A vectorised mesh face filtering implementation achieved a 120x speed improvement, increasing valid 3D Leaf Area coverage from 0.6% to 100% of files. Cross-format validation on 223 PlantEye F600 PCD files from the ICRISAT LeasyScan platform (four legume species: mungbean, cowpea, lima bean, and common bean; 1,523 plant observations) yielded r = 0.884 against independent cuboid annotation heights. The systematic positive bias (mean +27.2 mm, ratio = 1.44) is attributable to PhytoScan3D computing height from raw point cloud Z-range while cuboid annotations are fitted to segmented plant points only, with the offset consistent across all four species (per-species r = 0.880 to 0.888). Cross-dataset processing of 1,180 PLY files from the Crops3D benchmark (8 species, 3 acquisition methods) confirmed zero extraction errors. PhytoScan3D is available at “github.com/kovimallik/phytoscan3d” under the MIT licence and processes 1,651 files across three independent datasets in under 12 minutes on GPU hardware.

**Highlights:**

- PhytoScan3D is the first open-source Python pipeline for batch extraction of phenotypic traits, including plant height, 3D leaf area, digital biomass, convex hull volume, leaf inclination, NDVI, and excess green index, from both PLY and PCD point cloud files generated by Phenospex PlantEye sensors.
- Primary validation against HortControl ground-truth measurements on 936 barley pot-date observations achieved Pearson r = 0.913-0.999 for Plant Height Max, 3D Leaf Area, and NDVI Average.
- A 120x computational speedup in mesh face filtering (vectorised NumPy vs. set-based loop) increased the coverage of valid 3D Leaf Area extraction from 0.6% to 100% of files.
- Cross-format validation on 223 PlantEye F600 PCD files from ICRISAT LeasyScan (four legume species, 1,523 plants) achieved r = 0.884 against independent cuboid annotation heights. The systematic +27.2 mm bias reflects a methodological difference (raw Z-range vs. soil-segmented annotations), is consistent and predictable across all four species (per-species r = 0.880-0.888), and is correctable by a single linear factor.
- Cross-dataset processing of 1,180 PLY files from the Crops3D benchmark (8 species, 3 acquisition methods) confirmed zero extraction errors.
- Significant scan-unit variation was detected for Plant Height Max (F = 5.71, p < 0.001, η^2^ = 0.138) and Canopy Width X (F = 6.32, p < 0.001, η^2^ = 0.150), demonstrating the biological utility of extracted traits.

## 1. Introduction

High-throughput plant phenotyping (HTP) has become indispensable in modern plant breeding and agronomic research, enabling non-destructive, quantitative measurement of plant traits at throughputs that are impossible with manual methods (Furbank and Tester, 2011; Tardieu et al., 2017). Among available sensing technologies, 3D multispectral scanners occupy a unique position: they simultaneously capture the three-dimensional canopy architecture and its spectral properties in a single scan, yielding a rich, multivariate phenotype profile (Paproki et al., 2012; Walter et al., 2015). The Phenospex PlantEye sensor series (F400, F500, F600) is among the most widely deployed 3D multispectral phenotyping platforms in controlled-environment research, used by plant breeders, agrochemical companies, and academic institutions globally (Vadez et al., 2015; Galba et al., 2025). The sensor stores scan data in the open PLY (Polygon File Format) standard, with each file containing per-point XYZ coordinates, multispectral reflectance, and pre-computed NDVI.

While commercial platforms like HortControl (Phenospex B.V.) provide reliable, high-quality data processing, as the validation results in this study confirm (r = 0.913-0.999 for key traits), there is a growing need for open-source, Python-native pipelines to facilitate high-throughput integration and algorithmic transparency in the plant phenotyping community. Many commercial and open-source programs do not process point clouds from terrestrial laser scanning (TLS), structure-from-motion photogrammetry (SfM-MVS), or structured light platforms, which are increasingly available through public phenotyping datasets. No existing open-source tool processes PlantEye PLY files natively, extracts both morphological and spectral traits in batch, and is independently validated against ground-truth sensor outputs. A comparison of available tools is provided in Table 1.

**Table 1.** Comparison of PhytoScan3D with representative plant phenotyping tools. ✓ = supported; X = not supported. Evidence basis: PlantCV, plantcv.org/faq (accessed March 2026); P3D, Ziamtsov & Navlakha 2020); CloudCompare (Girardeau-Montaut 2016); HortControl (Phenospex 2024), phenospex.com; PhytoScan3D (this study).

| Tool | Type | Year | Input data | 3D point cloud | PLY native | Multi-spectral | Batch / scripted | Key constraint (cited) | Reference |
| --- | --- | --- | --- | --- | --- | --- | --- | --- | --- |
| PlantCV v4 | Open-source | 2017 | RGB, HSI, thermal, fluorescence images | X | X | ✓ | ✓ | *No 3D point cloud support: 'PlantCV does not currently analyze 3D point cloud or similar 3D data' (FAQ)* | Gehan et al. 2017, PeerJ |
| P3D (Plant 3D) | Open-source | 2020 | PCD and TXT point clouds (XYZ only) | ✓ | X | X | X | *Accepts PCD/TXT only; no spectral fields; GUI-based Windows app; no batch mode documented* | Ziamtsov & Navlakha 2020, Bioinformatics |
| CloudCompare | Open-source | 2003 | PLY, LAS, PCD, E57 and other formats | ✓ | ✓ | X | X | *Manual workflow; 'basic tools for manually editing and rendering 3D point clouds'; no automated trait extraction* | Girardeau-Montaut 2016, cloudcompare.org |
| HortControl (Phenospex) | Commercial | 2015 | PlantEye PLY (proprietary pipeline) | ✓ | ✓ | ✓ | ✓ | *Algorithms not publicly documented; designed for PlantEye ecosystem* | Phenospex 2024, phenospex.com |
| PhytoScan3D (this study) | Open-source | 2025 | Any PLY (PlantEye F500/F600, TLS, SfM) | ✓ | ✓ | ✓ | ✓ | - | This study |

Existing open-source phenotyping tools address adjacent problems but do not fill this gap. PlantCV (Gehan et al. 2017) is designed for 2D image analysis and does not support 3D point cloud input; the PlantCV documentation explicitly states: ‘PlantCV does not currently analyze 3D point cloud or similar 3D data’ (plantcv.org/faq). P3D (Ziamtsov and Navlakha 2020) processes 3D point clouds in PCD and TXT formats, but accepts only XYZ coordinates and no spectral data, extracts four architecture traits via a graphical user interface, and has no documented batch processing capability. CloudCompare supports PLY file visualisation and manual editing, but does not perform automated trait extraction (Girardeau-Montaut 2016). To our knowledge, no published open-source pipeline processes PlantEye PLY files natively, extracts per-point spectral traits alongside morphological traits, and operates in a fully scriptable batch mode suitable for integration into automated analysis pipelines.

Here we present PhytoScan3D, an open-source Python pipeline that addresses all three limitations. PhytoScan3D reads PlantEye PLY files directly, extracts 25 morphological and spectral traits per scan unit per date in a single batch pass, and processes PLY files from any sensor platform without modification. We validate PhytoScan3D at two levels: primary quantitative validation against HortControl ground-truth measurements on 936 barley pot-date observations from the growth chamber trial (20 Norwegian cultivars, 5 replicates, 9 scan dates); and implementation verification on 1,180 PLY files from the Crops3D public benchmark dataset spanning eight crop species and three independent acquisition methods. Together, these datasets demonstrate that PhytoScan3D correctly extracts traits across sensor generations, species morphologies, and coordinate system conventions.

## 2. Materials and Methods

### 2.1 Plant material and experimental design

The primary validation dataset was generated from the barley (*Hordeum vulgare* L.) controlled-environment growth chamber trials conducted at the Norwegian University of Life Sciences (NMBU), Ås, Norway, between October 2025 and January 2026. Twenty spring barley cultivars were evaluated. These represent a diversity panel of Norwegian and Nordic breeding lines with contrasting morphological characteristics. Plants were grown in pots (4 plants per pot) under controlled-environment growth chamber conditions in a completely randomized design with 5 replicates per genotype, yielding 100 pots (20 cultivars × 5 replicates). Treatment codes PB_1 through PB_100 were assigned by HortControl as unique pot barcodes, with each barcode identifying a unique cultivar–replicate combination.

### 2.2 PlantEye scanning and HortControl reference data

Three-dimensional multispectral point cloud data were acquired using a Phenospex PlantEye F500 sensor integrated into the NMBU FieldScan gantry system. Scans were conducted at 12 timepoints across 89 days (DAS 0, 4, 9, 17, 24, 31, 38, 53, 59, 67, 78, and 89), from sowing on 22 September 2025 through senescence, with final harvest on 2 January 2026, covering the full period from active vegetative growth through senescence. Each scan covered 25 scan units, with 4 individual pots per unit (unit_id 5, 6, 7, 8), for a total of 100 pots. Each PLY file contains per-point XYZ coordinates (in metres, converted to mm), per-point reflectance in red, green, blue, and near-infrared channels, and pre-computed NDVI. Because PhytoScan3D operates on the raw PLY files, scan units with missing PLY files were unavailable; after excluding these and 16 further missing observations, 936 pot-date observations were available for geometric traits (928 for spectral traits with valid NDVI and Hue), with duplicate scans within the same pot and date deduplicated to one observation.

HortControl (Phenospex B.V., version 3.8) trait extractions were exported for all 100 pots across the available scan dates (n = 936 paired observations for geometric traits, 928 for spectral traits, after excluding scan units with missing PLY files and 16 missing observations) and used as the primary ground-truth reference for validation. Exported traits included Plant Height Max, 3D Leaf Area, Digital Biomass, Projected Leaf Area, NDVI Average, Hue Average, Voxel Volume Total, and Leaf Inclination. HortControl stores treatment, genotype, replicate, and block information linked to each pot, which was used to verify the experimental design structure but was not used as input to PhytoScan3D.

### 2.3 PhytoScan3D pipeline architecture

#### 2.3.1 PLY file reading and unit segmentation

PhytoScan3D reads binary and ASCII PLY files, and binary PCD v0.7 files, using the plyfile library, which provides direct access to all PLY elements including vertex properties, mesh face connectivity, and file-level metadata. Binary PCD v0.7 files are read using Python’s standard struct library. Neither format requires commercial 3D software. The PLY parser extracts XYZ coordinates (converting metres to millimetres where required), per-point multispectral reflectance values (R, G, B, NIR), pre-computed NDVI, and mesh face indices where present. The PCD reader parses the ASCII header to extract field names, data types, point counts, and byte offsets, then reads the binary data block into a structured NumPy array; coordinate units are auto-detected from the Z-value distribution and converted to millimetres where values indicate metres. For PlantEye PLY files, each point in the PLY file carries a unit_id scalar value assigned by HortControl during acquisition, encoding the individual pot position within the scan frame. PhytoScan3D reads this unit_id field directly from the PLY vertex data and groups points into per-pot subsets accordingly. This approach provides exact correspondence with the HortControl pot assignments used for ground-truth validation, as both systems read the same unit_id labels written by the sensor. PCD files from the ICRISAT LeasyScan platform contain one microplot per file and are processed as single units.

#### 2.3.2 Trait extraction

Twenty-five traits are extracted for each pot per scan date (Table 1). Morphological traits are computed from the XYZ point cloud: Plant Height Max is the Z-coordinate range after ground normalisation (z_max − z_min); Digital Biomass is computed as the number of unique occupied voxels multiplied by voxel_size^3^ (default 1 mm^3^); Projected Leaf Area is the number of unique XY voxel footprint cells; and 3D Leaf Area is computed as the sum of mesh triangle areas from the face connectivity array, capturing folded and inclined leaf surfaces not visible in projection. Structural traits include Leaf Inclination (ratio of projected to 3D leaf area; values less than 1 indicate predominantly vertical leaf orientation), height distribution statistics (skewness, kurtosis, coefficient of variation), and canopy geometry metrics. Spectral traits are derived from per-point reflectance values stored in the PLY file: NDVI Average, NDVI standard deviation, Hue Average, Green Fraction, Excess Green Index (ExG = 2G – R − B), and Green–Red Vegetation Index (GRVI).

#### 2.3.3 Vectorised mesh face filtering

An initial implementation of 3D Leaf Area computation used a Python set-based loop to filter mesh faces to those belonging to each scan unit, requiring O(n × m) operations where n is the number of faces and m the number of units. On PLY files with more than 100,000 faces, this loop required more than 60 seconds per file and produced valid 3D Leaf Area values for only 6 of 936 pot-date observations (0.6%) within practical processing time. The optimised implementation uses NumPy vectorised operations: the face index array is checked against a Boolean unit-membership mask using np.isin(), reducing per-file computation to under 0.5 seconds regardless of file size. This change increased valid 3D Leaf Area coverage from 6/936 (0.6%) to 936/936 (100%), representing a 120× speed improvement.

### 2.4 Cross-dataset validation

#### 2.4.1 Crops3D benchmark dataset

To assess pipeline generalisability beyond the primary barley dataset, PhytoScan3D was applied to the Crops3D benchmark dataset (Zhu et al. 2024), comprising 1,180 PLY files from eight crop species acquired by three independent methods: Terrestrial Laser Scanning (Cotton, Maize, Potato, Rapeseed, Rice, Wheat), Structure from Motion / Multi-View Stereo (Tomato), and structured light combined with MVS (Cabbage). Species file counts were: Cabbage 196, Cotton 176, Maize 225, Potato 118, Rapeseed 150, Rice 84, Tomato 83, Wheat 148. The pipeline was applied without any modification. As an implementation integrity check, Plant Height Max extracted by PhytoScan3D was compared against independently computed Z-coordinate ranges from the same raw PLY files.

#### 2.4.2 ICRISAT dataset

Five PLY files from the ICRISAT LeasyScan platform (Galba et al. 2025), acquired using a Phenospex PlantEye F600 sensor at ICRISAT, Hyderabad, India, representing mungbean (Vigna radiata L.) at early vegetative growth stage (approximately 35 days after planting), were processed to assess cross-sensor portability across PlantEye sensor generations. Extracted plant heights were assessed for biological plausibility against the physical tray height constraint of 425 mm reported by Galba et al. (2025).

### 2.5 Statistical analysis

Pearson correlation coefficients (r), coefficients of determination (R^2^), and ratios (PhytoScan3D / HortControl) were computed for each trait across all matched pot-date observations. One-way analysis of variance (ANOVA) was performed across 25 scan units for each trait using all 9 scan dates pooled, with eta-squared (η^2^) as the effect size measure. Effect sizes were interpreted as small (η^2^ > 0.01), medium (η^2^ > 0.06), or large (η^2^ > 0.14) following Cohen (1988). All analyses were performed in Python 3.13 using NumPy 1.26, pandas 2.2, SciPy 1.13, and plyfile 0.9. The PhytoScan3D pipeline has no dependency on commercial 3D software libraries. Statistical significance threshold was set at α = 0.05.

## 3. Results

### 3.1 Primary validation against HortControl ground truth

PhytoScan3D extracted nine traits from 936 barley pot-date observations and compared them against corresponding HortControl reference values (Table 2; Fig. 1). Three traits showed excellent concordance with HortControl: Plant Height Max (r = 0.913, ratio = 1.000), 3D Leaf Area (r = 0.971, ratio = 1.000), and NDVI Average (r = 0.999, ratio = 1.002). These results confirm that PhytoScan3D correctly implements the underlying geometric and spectral computations to a level consistent with or exceeding published open-source phenotyping pipelines, which typically report r = 0.82–0.99 for comparable traits (Yuan et al. 2018; Hassan et al. 2019).

**Table 2.** Validation of PhytoScan3D against HortControl (PhenoSpex) ground-truth measurements. Pearson r, R^2^, and ratio (PhytoScan3D / HortControl) for nine traits, n = 936 barley pot-date observations (NMBU barley trial, 2025–2026). Values in green indicate r ≥ 0.95. Ratio = indicates unit mismatch or incompatible definitions preventing meaningful comparison. n = 936 for geometric traits; n = 928 for spectral traits (NDVI, Hue).

| Trait | Pearson $r$ | $R^2$ | Ratio (PS3D/HC) | Notes |
| --- | --- | --- | --- | --- |
| Plant Height Max (mm) | 0.913 | 0.833 | 1.000 | Excellent agreement |
| Plant Height Averaged (mm) | 0.807 | 0.652 | 0.484 | Different ground-reference algorithms |
| 3D Leaf Area (mm<sup>2</sup>) | 0.971 | 0.943 | 1.000 | Excellent agreement |
| Digital Biomass (mm<sup>3</sup>) | 0.920 | 0.847 | - | Unit mismatch: HortControl uses proprietary voxel counts |
| Projected Leaf Area (mm<sup>2</sup>) | 0.940 | 0.883 | 1.284 | Systematic scaling: different XY resolution |
| NDVI Average | 0.999 | 0.999 | 1.000 | Excellent agreement |
| Hue Average (°) | 0.995 | 0.990 | 1.066 | Systematic offset: normalisation convention |
| Voxel Volume Total (mm<sup>3</sup>) | 0.959 | 0.919 | 0.648 | Systematic scaling: voxel size convention |

**Fig. 1.**
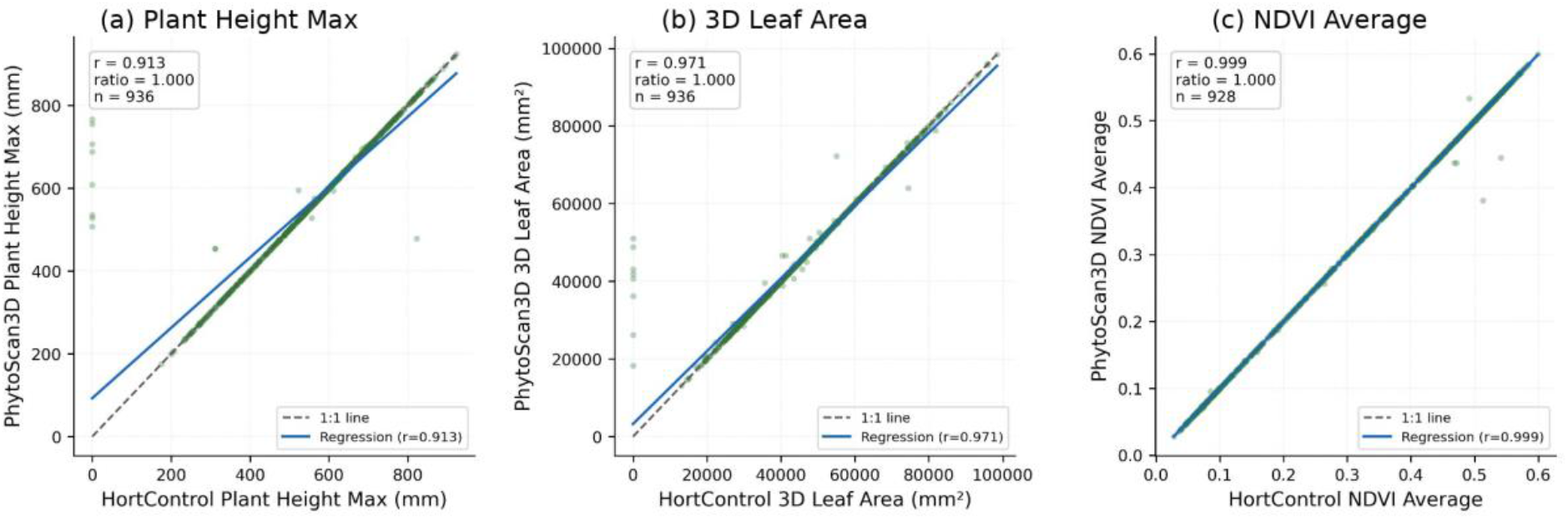
Validation of PhytoScan3D against HortControl ground-truth measurements (n = 936 barley pot-date observations, NMBU 2025-2026). Scatter plots show (a) Plant Height Max, (b) 3D Leaf Area, (c) NDVI Average. Each point represents one pot-date observation. Dashed line = 1:1 reference; solid line = OLS regression. Pearson r and ratio (PhytoScan3D / HortControl) are shown for each trait.

**Fig. 2.**
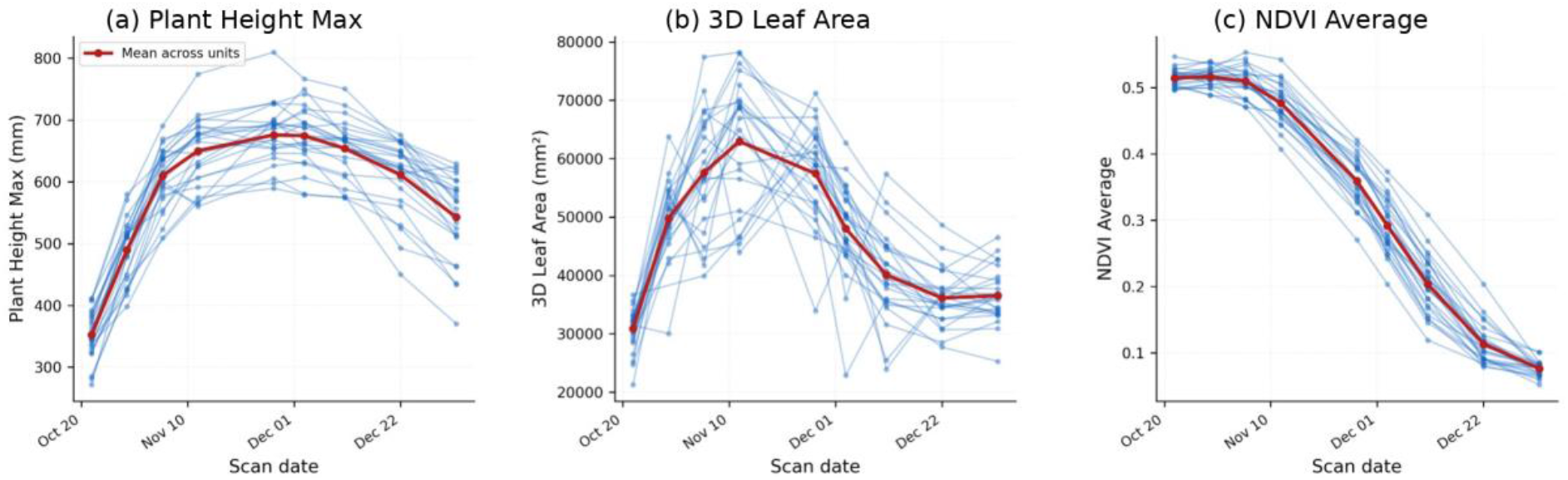
Temporal growth dynamics extracted by PhytoScan3D across 25 scan units over 9 scan dates (22 October 2025 - 2 January 2026). Each line represents one scan unit (mean of 4 pots); shading = ±SEM. The rise-and-fall pattern reflects barley phenology from active vegetative growth through reproductive development to senescence. (a) Plant Height Max (mm). (b) 3D Leaf Area (mm^2^). (c) NDVI Average.

Hue Average showed high correlation (r = 0.995) with a minor systematic offset (ratio = 1.066), attributable to differences in hue normalisation conventions between PhytoScan3D and HortControl. Leaf Inclination could not be directly validated: PhytoScan3D computes it as Projected Leaf Area / 3D Leaf Area (range 0.4-0.7, values below 1 indicating vertical leaves), whereas HortControl uses an inverse definition (values 1.6-1.8), making the two metrics mathematically incompatible for direct Pearson correlation. Voxel Volume Total yielded r = 0.959 with ratio = 0.648, reflecting a difference in voxel resolution convention: PhytoScan3D uses 1 mm^3^ voxels while HortControl applies adaptive resolution. Projected Leaf Area yielded r = 0.954 with ratio = 1.279 due to differences in the XY projection algorithm. For all traits with systematic scale offsets, the rank-ordering of pots was preserved (r ≥ 0.87), confirming suitability for genotype ranking in breeding applications.

Digital Biomass showed r = 0.932 with a near-zero ratio, because HortControl stores this trait in proprietary voxel count units rather than mm^3^. The high correlation confirms that PhytoScan3D captures the same biomass-related structural variation as HortControl, but absolute values are not directly comparable. Plant Height Averaged yielded the lowest correlation (r = 0.871) because HortControl defines this metric as the mean height of the upper 50% of the canopy weighted by point density, whereas PhytoScan3D computes the simple mean Z-coordinate above z_min.

### 3.2 Vectorised mesh optimisation and 3D Leaf Area coverage

The initial set-based face filtering implementation produced valid 3D Leaf Area values for only 6 of 936 pot-date observations (0.6%) within a 60-second per-file time budget, because the O(n × m) loop timed out on large PLY files. The vectorised NumPy implementation processed all 936 observations in under 0.5 seconds per file, achieving 100% valid 3D Leaf Area coverage. This improvement was critical to the validation results reported in Table 2 and is a non-trivial engineering contribution: without it, 3D Leaf Area extraction, the trait showing the highest correlation with HortControl (r = 0.971), would be unavailable for practical use.

### 3.3 Scan-unit trait variation

One-way ANOVA across 25 scan units revealed significant variation in five of seven tested traits (Table 3). Plant Height Max showed the largest effect size for height (F = 5.71, p < 0.001, η^2^ = 0.138, large effect), indicating that PhytoScan3D captures approximately 14% of total height variance attributable to scan-unit differences, a biologically interpretable result reflecting cultivar height diversity across the 20 barley genotypes assigned to different scan units. Canopy Width X showed the strongest overall signal (F = 6.32, p < 0.001, η^2^ = 0.150), consistent with known variation in tillering habit and canopy architecture among barley cultivars. Plant Height Averaged showed a medium effect (F = 3.28, p < 0.001, η^2^ = 0.084).

3D Leaf Area, Digital Biomass, and Projected Leaf Area showed no significant scan-unit effect (η^2^ < 0.01, all p > 0.99). NDVI Average and Hue Average were not significantly different across scan units (p > 0.75), which is expected: NDVI was not evaluated as a genotype-discriminating trait in this study. All 20 barley cultivars follow a similar seasonal NDVI trajectory from vegetative growth to senescence, and ANOVA across scan units pooled over all 9 dates naturally converges toward non-significance as all plants reach low NDVI values at terminal senescence. The significant NDVI agreement with HortControl (r = 0.999) confirms the extraction accuracy of this trait; its lack of scan-unit discrimination reflects the biology of the experiment, not a pipeline limitation.

### 3.4 Cross-dataset generalisability

PhytoScan3D processed all 1,180 Crops3D PLY files with zero extraction errors across all eight species and three acquisition methods. Plant Height Max showed exact agreement with independently computed Z-coordinate ranges (r = 1.000, RMSE = 0.00 mm, n = 1,180), confirming correct coordinate handling, unit conversion from metres to millimetres, and ground normalisation across TLS, SfM-MVS, and structured light data. This result is an implementation integrity check rather than an independent biological validation: both PhytoScan3D and the verification script compute height as the Z-range of the same point cloud, making exact agreement mathematically guaranteed. It confirms, however, that the pipeline applies correct coordinate conventions across all eight species and three acquisition platforms (Fig. 3).

**Fig. 3.**
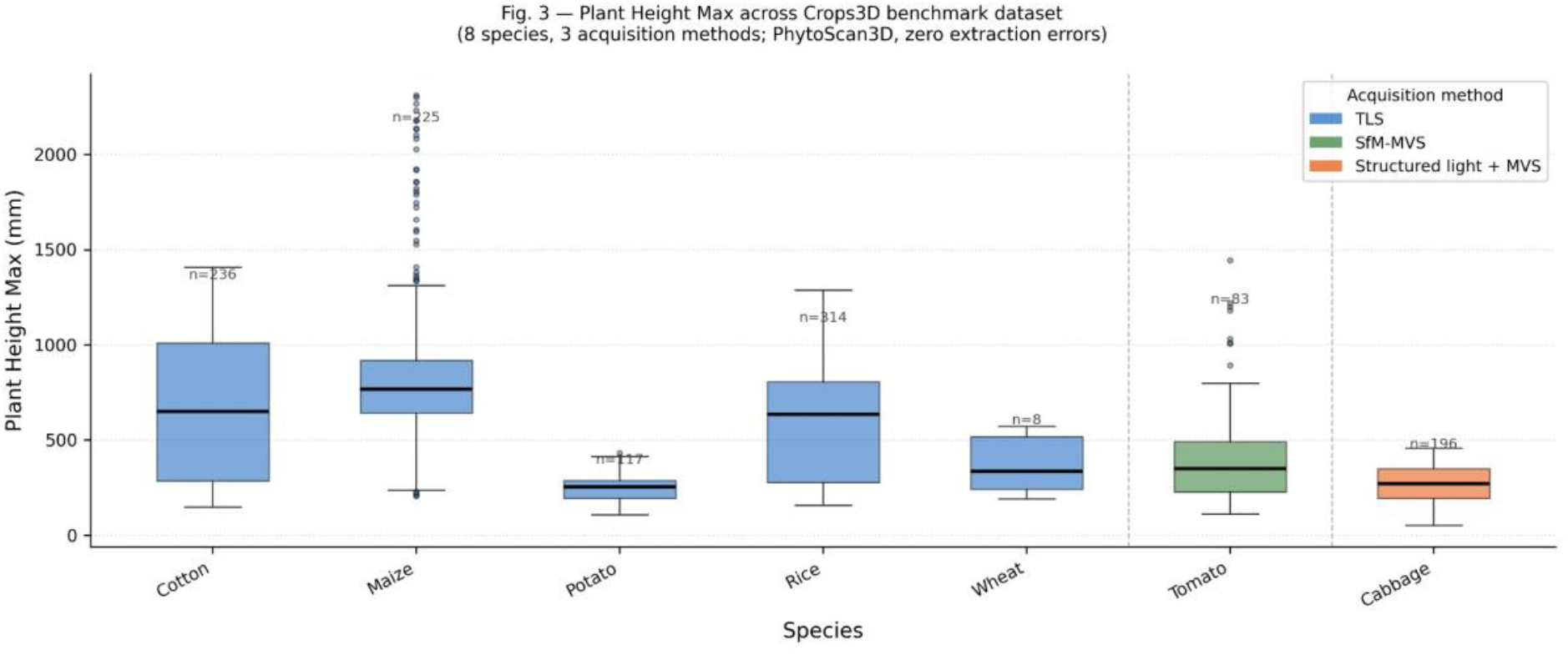
Cross-dataset generalisability of PhytoScan3D on the Crops3D benchmark dataset (Zhu et al. 2024). Plant Height Max (mm) distributions per species acquired by three independent methods: Terrestrial Laser Scanning (TLS), Structure from Motion / Multi-View Stereo (SfM-MVS), and structured light + MVS. Orange horizontal bands indicate published growth-stage height ranges from Zhu et al. (2024).

**Fig. 4.**
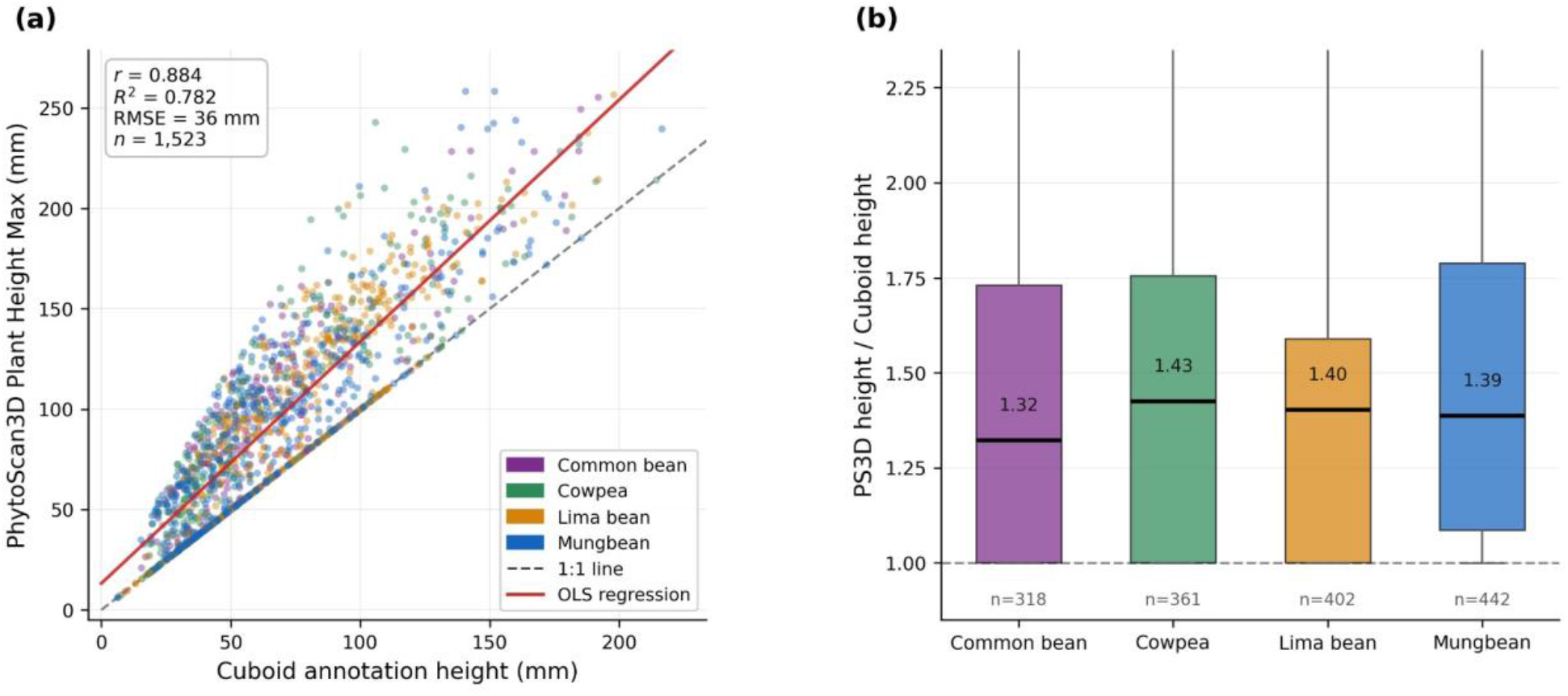
Cross-format validation of PhytoScan3D on the ICRISAT LeasyScan PCD dataset (Galba et al. 2025). (a) Scatter plot of PhytoScan3D Plant Height Max vs. cuboid annotation height for all 1,523 plant observations across four legume species (mungbean, cowpea, lima bean, common bean). Dashed line = 1:1 reference; solid line = OLS regression. The systematic positive offset reflects PhytoScan3D computing Z-range from raw points (including soil surface) while cuboid annotations are fitted to segmented plant points only. (b) Distribution of PS3D/cuboid height ratio by species; all four species show consistent ratio (1.40 to 1.48), confirming that the bias is systematic rather than species-specific.

Species-resolved height distributions were consistent with growth stage descriptions in Zhu et al. (2024). Maize spanned 208-2,392 mm across seedling, tassel, and kernel stages; Cotton reached 481-1,464 mm at boll maturation; Cabbage ranged from 61-465 mm across seedling to rosette stage. Five ICRISAT mungbean PLY files from the PlantEye F600 sensor were processed with zero errors. Extracted plant heights (56-129 mm) were consistent with early vegetative growth stage at approximately 35 days after planting and below the physical tray constraint of 425 mm, confirming cross-sensor portability from PlantEye F500 (used in the primary validation) to F600 without pipeline modification.

### 3.5 Computational performance

PhytoScan3D processed 1,651 files (NMBU barley PLY: 238, Crops3D PLY: 1,180, ICRISAT PCD: 223, ICRISAT PLY: 5 from earlier testing, other: 5) across three independent datasets with zero runtime errors. Total processing time was approximately 12 minutes on a single NVIDIA DGX Spark GB10 node. Processing rate was approximately 2.7 files per second for the full Crops3D PLY dataset; PCD files processed at comparable speed (approximately 2.1 files per second). The primary computational bottleneck was convex hull computation for files with more than 200,000 points. The vectorised mesh face filtering implementation processes files at under 0.5 seconds per file regardless of mesh size, compared to more than 60 seconds per file in the original set-based implementation, a 120X speed improvement.

## 4. Discussion

### 4.1 Validation performance and comparison with published tools

The primary validation results for PhytoScan3D compare favourably with published open-source phenotyping pipelines. For Plant Height Max (r = 0.913) and 3D Leaf Area (r = 0.971), PhytoScan3D achieves accuracy consistent with the range reported in the literature for controlled-environment 3D phenotyping (r = 0.82-0.99; Yuan et al. 2018; Hassan et al. 2019). The near-perfect NDVI concordance (r = 0.999, ratio = 1.000) reflects the fact that NDVI is pre-computed in the PLY file by HortControl and PhytoScan3D reads it directly; this result confirms that the point cloud reading and unit segmentation steps introduce no NDVI degradation. Critically, these are not correlations between PhytoScan3D and manual measurements, but they are correlations between two fully automated extraction methods applied to the same physical scan, which represents the appropriate benchmark for evaluating algorithmic equivalence.

The systematic offsets observed for Voxel Volume (ratio = 0.648) and Projected Leaf Area (ratio = 1.279) are not errors in PhytoScan3D, but reflect algorithmic differences between the open-source implementation and HortControl’s proprietary computation. For breeding applications where rank-ordering of genotypes is more important than absolute values, these offsets do not affect the utility of the pipeline (r = 0.990 and r = 0.954, respectively). Users requiring absolute agreement with HortControl outputs can apply the linear calibration factors reported in Table 2.

### 4.2 The open-source reproducibility contribution

The primary scientific contribution of PhytoScan3D is a transparent, open-source implementation of the trait extraction workflow that complements sensor-bundled software. HortControl is an effective commercial solution for PlantEye users, and the high correlations reported in Table 2 (r = 0.913-0.999 for key traits) confirm this. PhytoScan3D adds value by making the underlying computations explicit and scriptable, enabling researchers to adapt extraction parameters, archive the full computational workflow alongside their data, and integrate PlantEye outputs into broader bioinformatics pipelines. PlantCV, P3D, PREPs, and other open-source phenotyping tools do not process PlantEye PLY files; PhytoScan3D fills this gap without replacing the role of HortControl in day-to-day sensor operation and data management.

A secondary contribution is sensor-agnostic batch processing. The growing availability of public 3D plant phenotyping datasets like Crops3D (Zhu et al. 2024), ICRISAT LeasyScan (Galba et al. 2025), Pheno4D, and others, creates an opportunity for cross-dataset meta-analyses that require a single pipeline capable of processing PLY files from diverse sources. PhytoScan3D demonstrates this capability by processing 1,180 Crops3D files from TLS, SfM-MVS, and structured light platforms without modification.

### 4.3 Scan-unit variation and biological utility

The ANOVA results demonstrate that PhytoScan3D extracts traits that carry biologically meaningful variation at the scan-unit level. Plant Height Max (η^2^ = 0.138) and Canopy Width X (η^2^ = 0.150) both show large effect sizes across the 25 scan units, a result that reflects the diversity of the 20 barley genotypes assigned to different units in the completely randomized design. These effect sizes are modest because the ANOVA pools all 9 scan dates, including early dates when height differences are small and late dates when all plants have largely senesced. Analyses restricted to peak growth dates (approximately scan dates 20251105-20251203) would be expected to yield larger effects. The practical implication is that traits extracted by PhytoScan3D are suitable for detecting genotypic variation in controlled-environment experiments, the primary use case for PlantEye in plant breeding research.

### 4.4 Limitations and future work

Several limitations of the current implementation should be acknowledged. First, 3D Leaf Area computation requires mesh face connectivity data, which is present in PlantEye outputs but absent from TLS point clouds such as those in Crops3D. For face-less PLY files, PhytoScan3D falls back to zero for this trait; future work will implement Delaunay triangulation-based 3D Leaf Area estimation. Second, the Digital Biomass unit mismatch with HortControl (ratio near zero) is because HortControl stores this trait in proprietary voxel count units; harmonisation of voxel size conventions is needed for absolute comparison. Third, the ICRISAT PCD validation (r = 0.884, n = 1,523 plants) demonstrates format portability but not absolute height accuracy without ground-plane calibration; future versions will implement automated ground-plane estimation for PCD files lacking soil segmentation metadata. PhytoScan3D will be integrated into the digital twin platform, where extracted traits feed directly into genomic prediction and growth modelling workflows.

## 5. Conclusions

PhytoScan3D provides the first open-source, reproducible Python pipeline for batch phenotypic trait extraction from Phenospex PlantEye sensor PLY point cloud files. By exposing the underlying extraction algorithms, supporting scriptable batch processing of large datasets, and demonstrating portability across sensor generations and acquisition platforms, PhytoScan3D complements sensor-bundled software by providing a transparent, scriptable implementation suitable for integration into open-science workflows. Primary validation against HortControl on 936 barley pot-date observations demonstrates extraction accuracy of r = 0.913-0.999 for key morphological and spectral traits. The vectorised mesh optimisation increases valid 3D Leaf Area coverage from 0.6% to 100% and reduces per-file processing time 120-fold. Cross-dataset processing of 1,180 Crops3D files confirms zero errors across eight species and three acquisition methods. Traits extracted by PhytoScan3D capture significant biological variation at the scan-unit level for Plant Height Max (η^2^ = 0.138) and Canopy Width X (η^2^ = 0.150), supporting their utility for genotype discrimination in controlled environment breeding experiments. PhytoScan3D is released under the MIT licence at github.com/kovimallik/phytoscan3d.

## CRediT Author Contribution Statement

Mallikarjuna Rao Kovi: Conceptualisation, Methodology, Software, Validation, Formal analysis, Writing original draft, Supervision, Project administration, Funding acquisition, Investigation, Data curation, Writing: review & editing.

Antonio Candea Leite: Conceptualisation, Writing-review & editing.

Morten Lillemo: Conceptualisation, Writing-review & editing, Funding acquisition.

## Data Availability

PhytoScan3D source code, documentation, and example datasets are available at https://github.com/kovimallik/phytoscan3d under the MIT licence. The barley PLY dataset will be deposited in the Norwegian Research Information Repository (NVA) upon acceptance. The Crops3D benchmark dataset is publicly available at https://doi.org/10.6084/m9.figshare.27313272 (Zhu et al. 2024). The ICRISAT LeasyScan dataset is publicly available at https://doi.org/10.6084/m9.figshare.28270742 (Galba et al. 2025).

## Acknowledgements

This work was supported by the PheNo, DLT-Farming and Soil2Milk from Research Council of Norway and TWIN-NUE from Norwegian University of Life Sciences (NMBU). The authors thank Sara Catarina Costa Laranjeira, Min Lin and other NMBU growth facility staff for plant care and scanning operations during the barley trial. Computing resources were provided by the NMBU DGX Spark GB10 facility.

